# On the prevalence of uninformative parameters in statistical models applying model selection in applied ecology

**DOI:** 10.1101/448191

**Authors:** Shawn J. Leroux

## Abstract

Research in applied ecology provides scientific evidence to guide conservation policy and management. Applied ecology is becoming increasingly quantitative and model selection via information criteria has become a common statistical modeling approach. Unfortunately, parameters that contain little to no useful information are commonly presented and interpreted as important in applied ecology. I review the concept of an uninformative parameter in model selection using information criteria and perform a literature review to measure the prevalence of uninformative parameters in model selection studies applying Akaike’s Information Criterion (AIC) in 2014 in four of the top journals in applied ecology *(Biological Conservation*, *Conservation Biology*, *Ecological Applications*, *Journal of Applied Ecology).* Twenty-one percent of studies I reviewed applied AIC metrics. Many (31.5 %) of the studies applying AIC metrics in the four applied ecology journals I reviewed had or were very likely to have uninformative parameters in a model set. In addition, more than 40 % of studies reviewed had insufficient information to assess the presence or absence of uninformative parameters in a model set. Given the prevalence of studies likely to have uninformative parameters or with insufficient information to assess parameter status (71.5 %), I surmise that much of the policy recommendations based on applied ecology research may not be supported by the data analysis. I provide warning signals and a decision tree to help reduce the prevalence of uninformative parameters in studies applying model selection with information criteria. The four warning signals and decision tree should assist authors, reviewers, and editors to screen for uninformative parameters in studies applying model selection with information criteria. In the end, careful thinking at every step of the scientific process and greater reporting standards are required to detect uninformative parameters in studies adopting an information criteria approach.

## Introduction

Conservation biology emerged as a crisis discipline in the 1970s in response to evidence of widespread declines in biodiversity [1]. Along with the evolution of new technologies (e.g. Remote Sensing, Geographic Information Systems) and increasing availability of environmental (e.g. Land-use) and biodiversity (e.g. species occurrence records) data, the discipline has developed into a rigorous quantitative science [2-4]. These advances in methods and data allow applied ecologists to tackle complex problems at larger temporal and spatial scales than before. The application of quantitative analyses and the interpretation of these analyses in applied ecology is particularly important as research in this field often informs policy and management practices [5-7].

Around the same time as the field of conservation biology was emerging, Akaike [8] was paving the way for the broad application of the information criteria (IC) approach to statistics for evaluating data-based evidence for multiple working hypotheses [9, 10]. Model selection using IC is now a common type of analysis in applied ecology (Fig 1). This statistical approach encourages *a priori* development of multiple working hypotheses and presents formal methods for weighing the evidence supporting the different hypotheses [see reviews in 10-12]. As with any quantitative method, there are many challenges and ways to misuse IC techniques and recent work has highlighted some important issues in the application of IC in ecology, evolution, wildlife management and conservation biology. For example, Galipaud et al. [13] show how model averaging using the sum of model weights can overestimate parameter importance and Mac Nally et al. [14] present a plea for including absolute measures of model goodness-of-fit when possible as the top ranked model determined by IC may not be a “good” model. In addition, several researchers have called on the need for independent model validation [14-16] and better reporting of methods and results to facilitate critical evaluation of research conclusions (i.e. greater transparency [17]). Here, I focus on one issue in IC; uninformative parameters (*sensu* [18]). Uninformative parameters have received some attention in the literature [e.g. 11, 18-20] but this issue is still prevalent in applied ecology.

**Figure 1.**
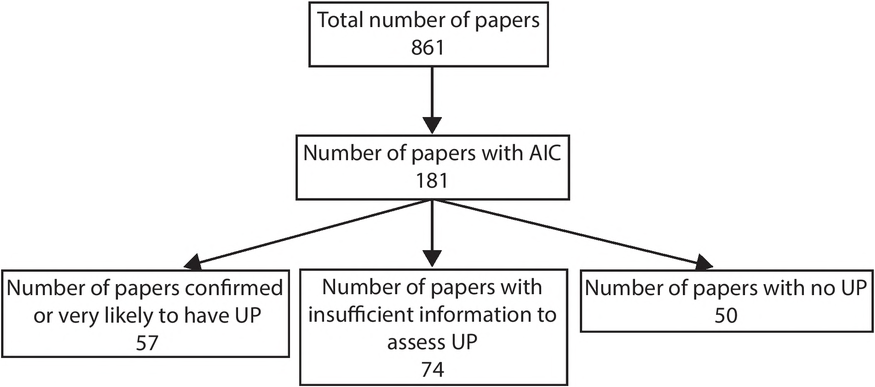
Summary of the use of IC and prevalence of uninformative parameters in articles reviewed from four top applied ecology journals *(Biological Conservation, Conservation Biology, Ecological Applications, Journal of Applied Ecology).* Articles were classified into four different categories for the prevalence of uninformative parameters in model sets – see main text for description of categories. Note that many papers in these journals do not use statistical analyses (e.g. essays). UP = uninformative parameter.

An “uninformative parameter” (*sensu* [18]) also known as a “pretending variable” (*sensu*[9, 11]), is a variable that has no relationship with the response, makes little to no improvement in the log-likelihood of a model (i.e. model fit) but can be included in a model ranked close to models with informative parameters based on IC. Interpreting uninformative parameters as important is a Type I error in statistics (i.e. false-positive [21]). If the interpretation of uninformative parameters as important is common, particularly in policy and management related fields such as conservation biology or medicine, then the policy recommendations of research may not be supported by the data analysis. What is more, poor data analysis and interpretation can lead to the natural selection of bad science (*sensu* [22]). My objectives are twofold; i) review and operationalize the concept of an uninformative parameter in model selection using IC and ii) quantify the prevalence of uninformative parameters in model selection using IC in applied ecology. Based on the results, I end with recommendations for how to screen for uninformative parameters in model selection studies using IC.

### Identifying uninformative parameters

In this section, I provide background on model selection using IC and formally present the concept of an uninformative parameter in this context. A clear definition of this concept is essential before presenting the methods and results of the quantitative review of uninformative parameters in model selection using IC in applied ecology.

Several IC exist for assessing the weight the evidence in support of different hypotheses formulated as competing statistical models [9] but I focus on the most commonly applied tool, Akaike’s Information Criterion (AIC) and related variations (e.g. AIC_c_). AIC is defined as

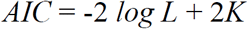

where *L* is the likelihood of the model given the data and *K* is the number of estimated parameters in the model. *K* is included as a penalty for adding additional parameters to the model, therefore AIC prioritizes parsimonious models. It is customary practice in model selection with IC to rank competing models from lowest to highest AIC or more specifically to rank models based on the △AIC which is the difference in AIC between a focal model and the model with the lowest AIC [9]. Top ranked models are models with a △AIC = 0. Models with △AIC < 2 are often considered equally supported or not differentiable from the top ranked model [9]. It is easy to see that two models (subscript 1 and 2) having identical log *L* (i.e. same fit to the data) but differing only by 1 estimated parameter (i.e. *K*_1_ − *K*_2_ = 1) will have a difference in AIC = 2. Given identical log L, the model with the additional parameter will have a *larger* AIC than the model with one less parameter and therefore the model with one additional parameter will be ranked *below* the simpler model. Likewise, models with identical log *L* and differing by 2 estimated parameters will have a difference in AIC = 4, and so on.

Uninformative parameters occur when there are nested models or more specifically, more complex versions of simpler models, in a model set [11, 18, 23-25]. Importantly, if the log *L* of a model has not improved with the addition of a parameter, it is likely that this additional parameter does not improve model fit and should be considered an uninformative parameter. However, if adding a parameter to a model improves the model fit, then the log *L* will increase and the AIC will decrease (i.e. the model with the extra parameter will be ranked *above* the model with one less parameter). See Table 1 for an illustration of uninformative parameters from some recent empirical research. Next, I outline the warning signals for uninformative parameters and summarize these warning signs in a decision tree that can be used to formally identify uninformative parameters in IC analyses (Fig 2).

**Table 1.** Here, I illustrate uninformative parameters from a real example derived from analyses in Yalcin and Leroux [26]. The objective of this study was to assess the relative and combined effects of land-use change and climate change on the colonization and extinction of species. We used a case study in Ontario, Canada where birds were surveyed in standardized grids during two time periods (1981-1985 and 2001-2005). Below I provide results for a subset of the colonization models of one of the study species, black-throated blue warbler (*Setophaga caerulescens).* In the colonization model, the black-throated blue warbler is observed as absent in a grid in the first time period and the response is warbler absence (0) or presence (1) in the second time period. We selected covariates based on *a priori* hypotheses. These covariates measured changes in land-use (% change in land-cover in each grid (%LCC), *%*change in land-cover in 20km buffers surrounding each grid (%LCCb) and change in Net Primary Productivity (ΔNPP)) and climate (change in mean winter temperature (ΔMWT), change in mean summer temperature (ΔMST), and change in mean winter precipitation (△MWP)) during the time period between bird sampling. All models include sampling effort (SE) in order to control for variable sampling effort across grids and between time periods. Yalcin and Leroux [26] fit generalized linear models with a binomial error structure and a logit link for local colonization models for the black-throated blue warbler. See for full details on data, methods, and hypotheses pertaining to each covariate used in these models. Table 1 provides a summary of AIC model selection results and parameter estimates (95% Confidence Interval) for a sub-set of the colonization models considered for this species. By following the decision tree in Fig 2, Yalcin and Leroux [26] identified the variable %LCCb is an uninformative parameter in models 2, 4, and 6.

| Model | SE | ΔNPP | %LCC | %LCCb | ΔMST | ΔMWT | ΔMWP | K | log L | ΔAIC<sub>C</sub> | *Pseudo R<sup>2</sup> |
| --- | --- | --- | --- | --- | --- | --- | --- | --- | --- | --- | --- |
| 1 | 0.30 | 0.06 | 0.04 |  | 0.25 | -2.54 |  | 6 | -276.14 | 0.00 | 0.28 |
|  | (0.24,0.39) | (0.03,0.09) | (0.02,0.06) |  | (0.10,0.40) | (-4.64,-0.46) |  |  |  |  |  |
| 2 | 0.30<br>(0.24,0.39) | 0.06<br>(0.03,0.09) | 0.04<br>(0.01,0.06) | **0.00**<br>**(-0.05,0.05)** | 0.25<br>(0.10,0.41) | -2.54<br>(-4.65,-0.47) |  | 7 | -276.14 | 2.00 | 0.28 |
| 3 | 0.30<br>(0.22,0.39) | 0.05<br>(0.02,0.08) |  |  | 0.25<br>(0.10,0.41) | -2.06<br>(-4.23,-0.09) | 0.07<br>(0.02,0.12) | 6 | -278.18 | 4.07 | 0.27 |
| 4 | 0.30<br>(0.22,0.38) | 0.05<br>(0.02,0.08) |  | **0.03**<br>**(-0.02,0.07)** | 0.29<br>(0.12,0.45) | -2.04<br>(-4.23,-0.12) | 0.07<br>(0.01,0.12) | 7 | -277.63 | 4.97 | 0.27 |
| 5 | 0.30<br>(0.22,0.38) | 0.06<br>(0.04,0.09) |  |  | 0.32<br>(0.18,0.46) |  | 0.08<br>(0.03,0.13) | 5 | -279.95 | 5.61 | 0.26 |
| 6 | 0.30<br>(0.22,0.38) | 0.06<br>(0.04,0.09) |  | **0.03**<br>**(-0.02,0.07)** | 0.35<br>(0.20,0.50) |  | 0.08<br>(0.03,0.13) | 6 | -279.34 | 6.39 | 0.26 |
| Intercept |  |  |  |  |  |  |  | 1 | -344.09 | 125.91 | 0.00 |

\*McFadden's pseudo R<sup>2</sup>

**Figure 2.**
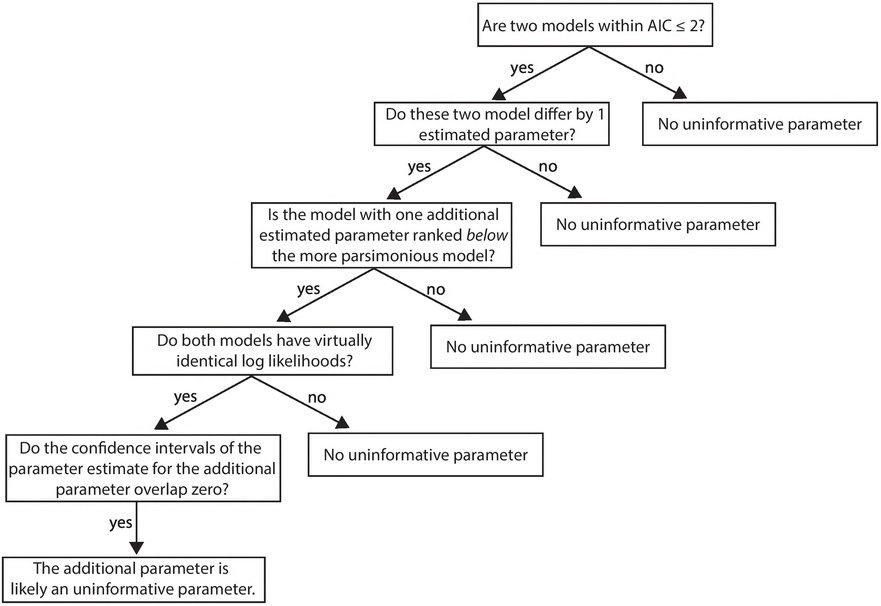
Decision tree for identifying models with uninformative parameters in a model set based on warning signals (see main text). This decision tree was used to assess the prevalence of uninformative parameters in top applied ecology journals (see Quantitative review). Note that the particular cut-off for the first step will vary based on the IC used (see main text).

Here I focus on cases where one additional estimated parameter may be an uninformative parameter but the logic also applies for cases where a model contains two additional estimated parameters and both may be uninformative parameters. These warning signals should be assessed in sequence (i.e. they build on each other, Fig 2). An uninformative parameter may exist in a model set if:

1. there are two models that differ by one estimated parameter that are within AIC ≤ 2 of each other. Authors must screen *all* possible model pairs in a model set (i.e. not just top ranked models) as a parameter may not be uninformative in every model in which it ^1^ appears given varying levels of multi-collinearity among covariates. Note that different IC metrics will yield slightly different cut-off points for detecting this first warning signal. For example, based on 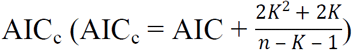 and a sample size (*n)* of 30, two models with identical log *L* and differing by only 1 parameter will have △AIC_c_ = 2.21. Consequently, the particular cut-off for this first warning signal should be considered in light of the specific IC metric used.
2. the model with one additional parameter (as outlined in warning signal 1) is ranked *below* the model with one less parameter (i.e. less parsimonious model AIC > more parsimonious model AIC). This suggests that the model with one additional parameter does not have a much better fit (i.e. log *L*) than the simpler model.
3. the models identified in warning signals 1 and 2 have virtually identical log *L.* Nearly identical log *L* suggests that the additional parameter is not contributing to improving model fit. This warning signal is subjective as there will be very few cases where the log *L* of two different models are identical. Consequently, authors must decide what is a sufficient difference to demonstrate that the added parameter contains useful information about the data. A strength of model selection with IC is that it allows researchers to use all available information to draw inference [9]. If authors are too strict in the cut-off for what they consider useful information, then authors risk losing inferential power. To avoid committing a Type I error, it may be best to err on the side of caution and to lose some information than to mis-interpret uninformative parameters as useful information. Given log *L* is a relative measure based on the data, there is no specific cut-off to determine if log *L* are similar. In lieu of a specific cut-off, researchers should assess parameter estimates and confidence intervals as a final step to identify uninformative parameters (see warning signal 4 [18]).
4. the additional parameter identified from warning signals 1-3 has a parameter estimate near zero with a confidence interval overlapping 0 [11, 18, 20, 27]. A parameter estimate near zero suggests that there is no relationship between this variable and the response variable. Arnold [18] and Galipaud et al. [20] provide specific guidance on confidence interval interpretations for identifying uninformative parameters.

By sequentially searching for the above warning signals, authors can identify all possible uninformative parameters in a model set (Fig 2). In order for readers of scientific papers to independently assess these warning signals, it follows that authors must provide all information required to interpret model selection with IC analyses.

While some recent research has demonstrated issues with uninformative parameters usually as part of broader studies [11, 18, 20, 25, 27], none have documented the prevalence of uninformative parameters in applied ecology and focused on solutions. Next, I provide a quantitative review of the prevalence of uninformative parameters in four of the top journals in applied ecology.

## Methods

I reviewed all 2014 articles in four of the top journals in applied ecology; *Biological Conservation, Conservation Biology, Ecological Applications*, and *Journal of Applied Ecology* for evidence of uninformative parameters. Specifically, I downloaded every article for each journal and I searched for the terms AIC or Akaike Information Criterion. I retained all articles with the term AIC in it. Following this first pass, I removed all articles that did not apply AIC in their analysis (i.e. they just mention AIC in the text).

I determined the presence or absence of uninformative parameters by systematically searching for the four warning signals in the order listed in the previous section and outlined in the decision tree (Fig 2). For warning signal 1, I only focused on pairs of models that differ by AIC ~ 2 and one estimated parameter. I used AIC ~ 2 as a cut-off as different articles used different AIC metrics (e.g. AIC, AIC_c_, qAIC). I did not focus on cases where two models differ by 2 or more parameters (i.e. differ by AIC ~ 4) – so my assessment of the prevalence of uninformative parameters is a minimum or conservative estimate. In many cases, authors did not provide sufficient information to fully determine if a model set included a model with an uninformative parameter. For example, AIC tables or estimates of model coefficients were often absent and when AIC tables were provided, key information such as the number of estimated parameters (*K*) or log *L* were often omitted. Consequently, I identified four possible uninformative parameter outcomes for each article in the study; i) articles with uninformative parameters, ii) articles with no uninformative parameters, iii) articles *very likely* to have uninformative parameters, iv) articles with insufficient information to identify uninformative parameters. These possible outcomes can be interpreted as follows. An article was classed as outcome i) if it had all four warning signals and outcome ii) if it did not have one of the warning signals. I assumed that the occurrence of one model with one uninformative parameter was sufficient to classify an article as having uninformative parameters. In most cases where there was one model with confirmed or very likely uninformative parameters, there were many models with uninformative parameters in the model set. I do not, however, report on the number of uninformative parameters per article. An article was classified as outcome iii) if it had the first three warning signals and as outcome iv) if there was insufficient information to assess any of the warning signals.

The article classification followed a two-step process. In the first step, two reviewers with experience in model selection with IC (lead author and A. Tanner (MSc working with lead author)) independently placed each article into one of the four outcomes listed above. In step two, the lead author reviewed the independent responses and flagged any articles with disagreement between reviewers (n = 16 or 9 *%* of studies). Then the lead author re-read and re-assigned each article that had initial disagreement between reviewers. I extracted the following information from each article: basic article information (authors, title, journal, issue, pages), IC used (i.e. AIC, AIC_c_, qAIC), the presence or absence of △AIC, parameter estimates, model averaging, and stepwise IC and the uninformative parameter ranking (i.e. yes, no, very likely, insufficient information). All data are available online [28].

## Results

The literature review revealed 329, 187, 163, and 182 articles published in 2014 in *Biological Conservation, Conservation Biology, Ecological Applications*, and *Journal of Applied Ecology*, respectively (Table 2). From this total, there were 87 (26 %), 22 (12 %), 33 (20 %), 39 (21 %) articles from *Biological Conservation*, *Conservation Biology*, *Ecological Applications*, and *Journal of Applied Ecology*, respectively that used AIC metrics in their analysis (Table 2, Fig 1). While only 21 % of articles (n = 181 / 861) in these journals apply AIC, many papers in these journals do not use statistical analyses (e.g. essays).

**Table 2.** Summary statistics (number and percentage of articles) of uninformative parameter assessment for four top journals in applied ecology. Articles were classified into four different categories for the prevalence of uninformative parameters in model sets – see main text for description of categories. The number of articles and percent of articles reported are compared to the subset of articles with AIC per journal, except in the final row which reports the totals across all journals. UP = uninformative parameter.

| Journal (Total # in 2014) | Total #<br>(%) with<br>AIC | Number of articles (%) with |  |  |  |
| --- | --- | --- | --- | --- | --- |
|  |  | UP | very<br>likely UP | no UP | insufficient<br>information |
| Biological Conservation (329) | 87(26) | 7(8) | 20(23) | 25(29) | 35(40) |
| Conservation Biology (187) | 22(12) | 1(5) | 7(32) | 5(23) | 9(41) |
| Ecological Applications (163) | 33(20) | 0(0) | 8(24) | 8(24) | 17(51) |
| J. of Applied Ecology (182) | 39(21) | 3(8) | 11(28) | 12(31) | 13(33) |
| Total (861) | 181(21) | 11(6) | 46(25) | 50(28) | 74(41) |

Across all journals there was at least one model with an uninformative parameter in an article’s model set in 6 % of cases and no model with an uninformative parameter in an article’s model set in 28 % of cases. Only 4 % of articles self-identified uninformative parameters and removed them from their model set. *Biological Conservation* and *Journal of Applied Ecology* had the highest percentage of articles adopting an AIC approach where the presence or absence of uninformative parameters could be confirmed (Table 2). This statistic goes hand in hand with the fact that these two journals had the lowest percentage of articles with insufficient information to assess uninformative parameters, albeit these percentages were still high (*Biological Conservation* = 40 %, *Journal of Applied Ecology* = 33 %). *Ecological Applications* had no confirmed cases of models with uninformative parameters but it also had the highest percentage of articles with insufficient information to identify uninformative parameters (51 %, Table 2). Note that in many cases, there is no possibility for uninformative parameters as a model set may be very simple with a null model (i.e. intercept only) and one additional model with one fixed effect or a set of non-nested models (i.e. models with no overlapping parameters). For example, Barnes et al. [29]’s model set to investigate the response of dung beetle communities to land-use management in Afromontane rainforests in Nigeria included four non-nested models and therefore there is no possibility for uninformative parameters in their model set. Consequently, the percentage of studies with no uninformative parameters should be higher than the percentage of studies with uninformative parameters.

In 23 to 32 % (grand mean 25 %) of articles across the four journals there was evidence that uninformative parameters were very likely based on the information presented in the article (i.e. warning signals 1-3 were confirmed, Fig 1). Altogether, nearly 1/3 (31.5 %) of all articles considered had or were very likely to have models with an uninformative parameter in the model set (Table 2, Fig 1).

## Discussion

Applied ecologists are increasingly being called on to support evidence-based environmental and natural resource management. The evidence we provide, therefore, must be based on sound empirical design, statistical analyses, and interpretations of these analyses [5]. In this study, I conducted a quantitative review of the prevalence of uninformative parameters in model selection using IC in applied ecology. My review revealed two main findings with potential impacts on the field of applied ecology; i) many articles applying model selection with IC in this study had or were very likely to have at least one model in a model set with one uninformative parameter (Table 2, Fig 1) and ii) many articles had insufficient information to identify uninformative parameters in their model set. These two issues stand to reduce the validity of inference drawn from statistical analyses applying model selection using IC in applied ecology.

In many of the articles reviewed herein, uninformative parameters were reported as important and often interpreted as such. For example, *Biological Conservation* [30 – author names withheld] report the following results for two competing models (i.e. y ~ time; y ~ time + weather) of florican (*Sypheotides indicus)* detection in semiarid grasslands in India: “The time model had smallest AIC_c_ value, more precise effect (*β* = 0.62_Mean_ ± 0.31_SE_) and parsimony than the time and weather model (△AIC_c_ = 1.54, *β* = 0.56 ± 0.31 [time], 0.28 ± 0.34 [weather]). Time had stronger influence (AIC_c_−wt = 0.61) than weather (AIC_c_−wt = 0.31) on display frequency…”. The two models differ by one parameter, have almost identical log *L* (i.e. differ by 0.69) and the parameter estimate for weather overlaps zero. In this case, weather is an uninformative parameter and weather should be removed from the model set and presented as having little to no support (i.e. not interpreted as important). In contrast to this example many of the papers that did have uninformative parameters did not interpret these parameters as important. For example, Rudolphi et al. [31] have many uninformative parameters in their model sets to investigate the impacts of logging on bryophytes and lichens. However, they restrict their interpretation to parameters with 95*%* confidence estimates that do not overlap zero.

The quantitative review revealed that more than 40 % of all articles had insufficient information to identify uninformative parameters (Table 2, Fig 1). This lack of transparency in reporting of methods and results has been highlighted previously [e.g. 10, 17, 32, 33]. The missing information ranged from not reporting the number of parameters or log *L* per model, to not reporting parameter estimates, and in many cases not presenting any AIC table.

Based on my findings, I present the following recommendations for reducing erroneous interpretation of uninformative parameters from model selection studies in applied ecology. First, once authors have identified all uninformative parameters in a model set, I recommend that all models with uninformative parameters be removed from the model set and that the model removal be noted in the results section (see discussion of full reporting below; [11, 18]). In some cases, the top model may include an uninformative parameter uncovered elsewhere in the IC table and in such cases, the original top model should be removed from the model set. Models with interaction terms (i.e. X_1_ * X_2_) where a component (e.g. X_1_) of the interaction is an uninformative parameter in the model set should be retained because a parameter may be informative (i.e. improve model fit) once it is in interaction with another parameter. However, if an interaction term is an uninformative parameter, then all models with the full interaction term should be removed from the model set. The type of variable (i.e. continuous or categorical) will influence the approach to removing models with uninformative parameters. Continuous variables and categorical variables with two levels usually have one estimated parameter and advice for removal of uninformative parameters above can be followed. Categorical variables with more than 2 levels will have n – 1 estimated parameters where n is the number of levels. It is possible that one level of a multi-level categorical variable is uninformative but others are informative. In these cases, authors should retain the categorical variable but interpret the results for every level making a clear distinction between the informative and uninformative levels.

Second, a solution to detecting and removing uninformative parameters from analyses is to report sufficient information to assess the warning signals of uninformative parameters (see Fig 2, [11, 18]). Proper reporting of quantitative analyses should be a default in scientific research. Transparency will allow peer review to help identify uninformative parameters at various stages of the review process. At minimum, papers using model selection with IC must report AIC tables with *K*, log *L*, ΔAIC, absolute measure of goodness-of-fit (see [14]) and parameter estimates with some measure of confidence intervals for all models [9, 10]. Abbreviated AIC tables (i.e. models with △AIC < 8) may occur in the main text as per Burnham et al. [10] but the AIC table for the full model set prior to removal of models with uninformative parameters should be placed in supplement. Graphical presentations of modeled relationships also may be useful for understanding relationships [34, 35] and detecting uninformative parameters.

As described in Arnold [18], authors must not sacrifice full reporting when removing models with uninformative parameters. Specifically, authors should present all models in the methods and report the presence of uninformative parameters and subsequent model removal in the results. If done correctly, readers should be able to identify all models considered by authors and the particular parameters that were uninformative. Examples for clear reporting of all models considered and removal of uninformative parameters can be seen in Devries et al. [36], Fondell et al. [37], Beauchesne et al. [38] and Fitzherbert et al. [39].

Third, some IC techniques are more prone to uninformative parameters than others and steering away from such approaches can help reduce the occurrence of uninformative parameters. Cade [40] and Galipaud et al. [13, 20] convincingly demonstrate the perils of model averaging by summed IC weights (but see [41]). Most articles considered in the quantitative review which used model averaging by summed AIC weights were very likely to have uninformative parameters. For example, *Conservation Biology* [42–  author names withheld] present summed AIC weights for several models with uninformative parameters for the effects of land-use (i.e. mining vs agriculture) on West African rainforest bird richness (see their Figs 3 and 4).

Stepwise AIC runs counter to the original intention of model selection with IC [9-12, 21]. Stepwise AIC does not encourage the creation of *a priori* hypotheses and models but is rather usually applied to all possible models. Stepwise AIC was common in the studies reviewed with 14% of articles using some form of stepwise AIC in their analysis. The process of fitting all possible models without *a priori* reason is flawed [9-12, 21] and will often inflate the occurrence of uninformative parameters relative to an *a priori* model selection approach [27]. Note that uninformative parameters may still occur in a model set based on *a priori* selection of variables. However, trying all possible models will almost surely lead to more uninformative parameters. Stepwise AIC also does not allow one to assess model selection uncertainty [27] which is a critical component of multiple hypothesis testing. While stepwise AIC has critical flaws, the end result likely does not include uninformative parameters as the stepwise process ends with one top model and models with additional variables but higher AIC would have been thrown out during the stepwise process. That said, stepwise AIC should only be used when paired with *a priori* selection of variables and models.

Common advice to reduce uninformative parameters is to remove more complex or nested versions of simpler models in a model set [12, 24, 25]. This approach is not new to statistics [24] and it is commonly used in a Bayesian framework [43]. The articles in the data set that used this approach [e.g. 38, 39, 44] did not have uninformative parameters. Authors should think critically about nested models and only use the more complex versions of nested models if they represent *a priori* hypotheses for the phenomenon of interest.

## Conclusion

I provide quantitative evidence of the prevalence of uninformative parameters in IC studies in applied ecology and recommendations on how to diagnose and remove these uninformative parameters. My review focused on the most widely used IC metric; AIC, but uninformative parameters should be considered when applying other IC metrics (e.g. Bayesian Information Criterion, Deviance Information Criterion). Model selection with IC is a powerful tool to assess the evidence supporting multiple working hypotheses but only if the tool is applied correctly. Given the close connection of applied ecology to conservation policy and management, careful thinking at every step of the process from the individual researchers (i.e. study design, statistical analysis, interpretation of results), reviewers (i.e. interpretation of results, transparency in reporting), and editors is required for valid inferences to be made. Additional vigilance can be facilitated by improving the reporting standards for statistical analyses [35, 45] and by screening the statistical analyses of submitted articles. In the end, researchers must be critical of results and seek statistical advice when in doubt - biodiversity and the reputation of the field of applied ecology depends on it.

## Acknowledgements

I thank the A. Tanner for assistance with the literature review and S. Yalcin for providing data for Table 1. I am grateful to the Eco&Evo discussion group at Memorial University for constructive feedback. A. Buren, M. Laforge, K. Lewis, A. McLeod, C. Prokopenko, and Q. Webber read an earlier version of the ms and provided excellent suggestions which improved the content. This research was funded by a Discovery Grant from the Natural Sciences and Engineering Research Council of Canada.

## Data accessibility

Data available from figshare digital repository https://doi.org/10.6084/m9.figshare.6002582.v1

